# Nonlinear continuum description of the *E. coli* cell wall under high turgor pressure

**DOI:** 10.1101/2025.01.20.633943

**Authors:** Octavio Albarrán, Renata Garcés, Giacomo Po, Christoph F. Schmidt, Jeff D. Eldredge

**Affiliations:** Duke University, Department of Physics & Soft Matter Center, Durham, NC; University of California, Department of Mechanical & Aerospace Engineering, Los Angeles, CA; University of Miami, College of Engineering, Coral Gables, FL

## Abstract

The principal stress-bearing component of bacterial cell walls is the tough, covalently linked peptidoglycan network. It is an active material that maintains shape and integrity of bacterial cells in the presence of high turgor pressures while constantly growing. It is not well understood how its extremal material properties derive from its molecular structure. The large stress in the cell wall demands a description going beyond small-strain approximations, an approach that has not been taken in the biomaterials community. We here derive a continuum elastic model of *E. coli* peptidoglycan (PG) in two steps: We first reduce the complex microscopic structure of the PG layer to the spring elements of a 2D hyperelastic triangular network, and then interpret those results in the context of finite-strain continuum theory. We present a self-contained formalism to study stress-strain relationships for large deformations and arbitrary anisotropies. We obtain the minimal model compatible with the available data on *E. coli* morphology and mechanics. We show that the strain hardening of PG leads to a stiffness matrix that is linearly proportional to the turgor pressure of the cell.

---

Peptidoglycan networks are the layers in bacterial cell walls that bear the largest part of wall stress [1]. While the precise chemical composition and architecture of the peptidoglycan network (PG) vary between species, the PG is generally a porous polymer network made of rigid glycan strands cross-linked by short flexible oligopeptides [2]. Cell wall mechanics are crucial for bacterial morphogenesis and homeostasis [1, 3–6]. Cell walls are also the target of antibiotics of the beta-lactam family [7]. The understanding of the connection between the macroscopic mechanics of PG and its molecular constituents is incomplete.

In gram-negative *E. coli*, the PG network is about 1-3 molecular layers thick (2-7 nm) [8, 9]. The cell wall of *E. coli* has been probed with micro-manipulation techniques such as AFM [10, 11], microfluidics [12–14], geometrical confinement [15, 16] and osmotic shocks [17–20]. Continuum models used for data analysis typically rely on linear-response approximations valid for small strains. This is likely not the most appropriate approach since, under physiological conditions, *E. coli* maintain a substantial turgor pressure [17], which implies that the PG operates in the large deformation regime where non-linear responses are expected [13, 19–22].

Structural anisotropy is another crucial feature of PG. Glycan strands are more than two orders of magnitude stiffer than peptide crosslinkers [6] and it is commonly assumed that they are preferentially oriented in the direction of maximum stress, the hoop direction for rodshaped bacteria. In the absence of definitive data, highly ordered structural models with extreme anisotropies have been used to compute elastic constants [23, 24] and model the growth process [4–6, 25]. In experiments, *E. coli* sacculi were found to be about two times more deformable in the axial direction than in the hoop direction [8].

We here present a coarse-grained elastic model for PG that accounts for structural anisotropy and response non-linearities by mapping the mechanical properties into a 2D hyperelastic triangular network that can be treated analytically. In the continuum limit, a regular triangular network allows for the evaluation of general constitutive equations. We analyze the elastic responses in the context of finite-strain theory, valid for the large deformations observed in *E. coli* under physiological conditions. Hyperelastic materials are completely determined by their energy density. We construct this energy density using local information and derive all the mechanical measures, such as stress and stiffness, in the continuum limit of vanishing mesh size. The triangular network is a first step toward connecting the macroscopic properties of PG to its microscopic constituents, and has the advantage of being analytically treatable. In part I, we introduce a general continuum description of regular triangular 2D networks, valid for large deformations and arbitrary anisotropy. In the absence of damage or yield, materials typically stiffen under large strain. We show how, in general, the stiffening behavior of a material emerges from both, intrinsic strain hardening of its components and from geometric nonlinearities due to finite strain. We generalize the conventional stiffness measures to the large deformation regime. We then, in part II, construct a minimal model for *E. coli* PG based on experimental data on morphology and mechanics.

## I. HYPERELASTICITY OF REGULAR TRIANGULAR NETWORKS

Our aim in this first part is to describe the non-linear response of a regular triangular network and interpret this response as that of a hyperelastic continuum under finite (*i.e*. non-infinitesimal) strain. Finite-strain theory requires some care to (i) differentiate between the non-deformed and the deformed states of the material and to (ii) ensure that the material response is described objectively (*i.e*. does not depend on the frame of reference) [26]. In the following, we briefly review the important concepts to apply and the relevant quantities to define.

### A. Results from hyperelasticity

To connect the discrete network model with the continuum description, we define an elastic energy per unit resting area, 𝒲, of the network. A continuum material whose mechanics are derived from such an energy density is, by definition, hyperelastic [27]. In such a material, it is essential to distinguish between the resting configuration (for the bacterial cell wall this is the turgor-free state) and the configuration in the deformed state (*e.g*. under finite turgor pressure) because the definition of stress, for example, involves the choice of a surface that a force is acting on. The resting configuration serves as the basic reference for all turgid states. The parametrization of the deformed and resting configuration are related to each other via the deformation gradient tensor ***F***.

To characterize the material response in a deformed state, one can measure the changes in stress that result from a small additional strain applied to the already deformed turgid cell wall (pre-strained state). There are different possible definitions of stress in finite-strain theory. The most useful form of the stress is the one that we can “directly” measure, *i.e*. calculated from the forces and surfaces measured in the deformed state; this definition leads to the Cauchy stress tensor, ***σ***, also called the “true stress”, which is, in general, different from the stress calculated using the surface of the resting configuration.

The Cauchy stress is an objective measure of stress, which means that it transforms as expected if we change the frame of reference. This is not true, however, when we deal with differentials of Cauchy stresses. The stress difference δ***σ*** is *not* an objective measure because ***σ*** requires a transformation to conform with the frame of reference. If we were to naively measure the difference between Cauchy stresses when we deform the material by a small additional amount, we would fold in artifacts related to the change in observer perspective. This observer dependence poses difficulties when describing the material’s constitutive behavior, which must necessarily be independent of the frame of reference. To overcome this problem, finite strain theory typically gives up on using “true stress” and instead relies on deformation measures that use the surface of the resting configuration to compute nominal stresses. Two commonly used nominal tensors are the 2nd Piola-Kirchhoff stress tensor, ***S***, and the Green-Lagrange strain tensor,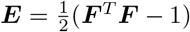.

The stress is connected to the energy density,𝒲, through the principle of virtual work, which in terms of the nominal tensors, ***S*** and ***E***, reduces to the simple form

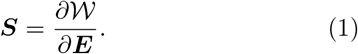

We note that here and throughout the rest of this paper, the energy density is defined per unit area, as is typical in two-dimensional materials, and therefore the stresses are expressed as units of force per unit length. Using ***S***, the objective relationship between small changes in stress and strain can be simply written as

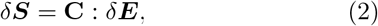

where δ denotes an incremental change, the colon symbol denotes a double dot product, and **C** is the rank-4 nominal elasticity tensor of the material,

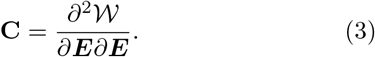

Moreover, the relationship between the Cauchy stress ***σ*** and ***S*** reads

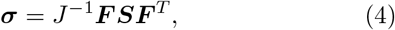

where *J* is the determinant of ***F*** and ()^*T*^ denotes transposition. To write an equation equivalent to (2), but involving the increment in Cauchy stress, we must re-define this increment so that it remains an objective measure of the stress change. One way to do this is via the Truesdell increment, δ_T_(*·*) [26], whose definition ensures that incremental changes in ***σ*** and ***S*** are related in the same way as ***σ*** and ***S*** themselves. With this definition, we get the relationship

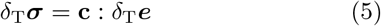

between the increments in Cauchy stress and the Euler-Almansi strain, ***e*** = ***F*** ^−*T*^ ***EF*** ^−1^, where **c** is the spatial elasticity tensor, whose components are related to those of the nominal elasticity tensor **C** by

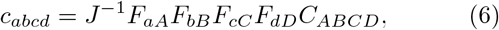

in which the summation convention of index notation has been used. For the nominal tensors, ***S*** and ***E***, the Truesdell increment is equivalent to the simple incremental change in the quantity. This is not so for ***σ*** and ***e***, and in particular, the Truesdell increment in Cauchy stress is

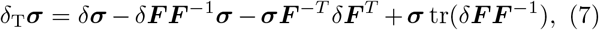

where tr denotes the trace. The Truesdell increment removes all of the non-objective effects from the measured stress increment, δ***σ***. The incremental deformation gradient tensor *δ****F F*** ^−1^ can be split into symmetric and skew-symmetric parts, *δ****F F*** ^−1^ = *δ****ε*** + *δ****ω***. The symmetric part, *δ****ε***, is the conventionally-defined infinitesimal strain, and can be shown to be equivalent to the Truesdell increment of ***e***, *i.e*. *δ****ε*** = *δ*_T_***e***. The skew-symmetric part represents rotational strain. Thus, the total change in Cauchy stress reduces to

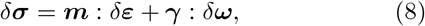

where, ***m*** is a modified version of the spatial elasticity tensor and ***γ*** is a rotational influence tensor [28]. In index notation:

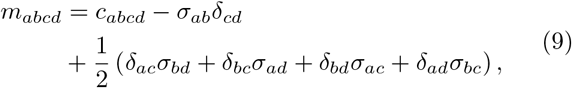

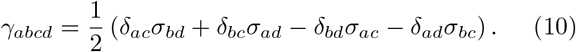

The components of ***m*** are analogous to the conventional elastic moduli for small deformations, while those of ***γ*** are non-conventional. From (8)-(10), it is apparent that the elastic response of the material to a small incremental deformation depends on its current state of stress. This is manifested by the explicit stress coupling of ***m*** and ***γ***, which comes from the observer dependence terms in (7), *δ****σ*** − *δ*_T_***σ***, and describe a geometric stress-stiffening behavior that is not associated with the intrinsic elasticity, *i.e*. it can occur in linear elastic materials. Even during a strain-softening response, where **C** decreases with increasing strain, the diagonal elements of ***m*** are non-zero and could increase with strain. We will discuss this geometric nonlinearity in the sections that follow. To build a bridge to a molecular description of the bacterial cell wall, we will use the energy density of a triangular network and relate its behavior to the continuum description of incremental strain applied to a pre-strained material.

### B. Regular 2D triangular networks

In 2D, the triangular unit cell of the network has three edges, each consisting of a spring. In the most general case, the three springs have different stretching rigidities and torque hinges, providing six independent stiffnesses, equal to the maximum number of independent elastic constant of a 2D material. We associate the lengths and orientations of the springs with three edge vectors, which are denoted by ***a***_1_, ***a***_2_, ***a***_3_ = ***a***_2_ − ***a***_1_, forming internal angles *θ*_1_, *θ*_2_, *θ*_3_ = *π* − *θ*_1_ − *θ*_2_ (FIG. 1). From hereon, the subscript *i* will denote the three springs (*i* = 1, 2, 3). Summation will be shown explicitly and is not implied by repeated indices. The values of any parameter or function of the network in the non-deformed state will be denoted by ()^*′*^, and those in the deformed state will be unprimed. The vectors of the resting and deformed configurations are related via the deformation gradient tensor ***F*** by

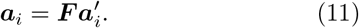

**FIG. 1:**
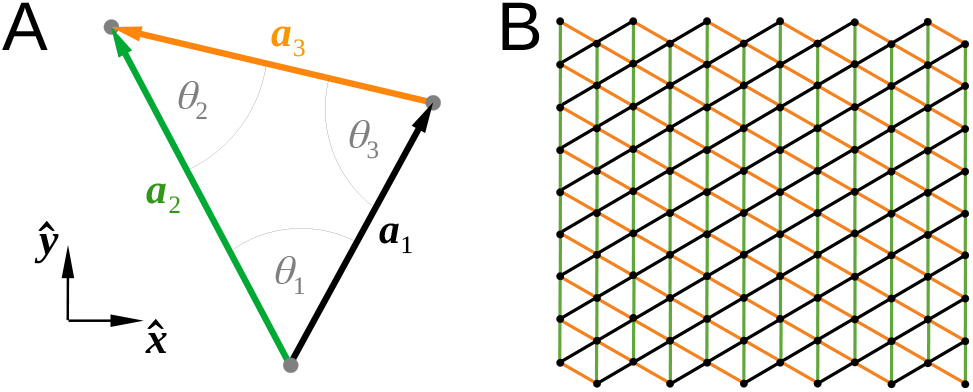
(A) Schematic of the unit triangle of a 2D regular network, with edge vectors ***a***_1_, ***a***_2_, ***a***_3_ = ***a***_2_ − ***a***_1_ and internal angles *θ*_1_, *θ*_2_, *θ*_3_ = *π* − *θ*_1_ − *θ*_2_. (B) Regular network patch generated with an equilateral triangular unit cell (hexagonal lattice).

In the regular 2D network, all springs of the same type will deform identically. The deformation of each type of spring can be expressed in terms of its intrinsic strain,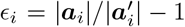.It follows that the unit vectors, ***â***_*i*_ = ***a***_*i*_*/* |***a***_*i*_| and 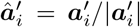, of the deformed and resting configurations, respectively, are related by

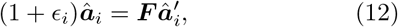

and furthermore,

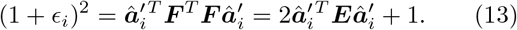

From this, we can derive the identities in (14) that will be useful for calculating the stress and stiffness through the energy-based relationships in (1) and (3):

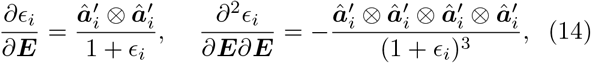

where ⊗ denotes the tensorial product.

The elastic energy density of the network (energy per unit of resting area) can, in principle, be any function of the form 𝒲 (*ϵ*_1_, *ϵ*_2_, *ϵ*_3_). In the continuum limit, since all springs in the network are shared by two triangles, 𝒲 is half of the energy density of the isolated unit triangle. Our goal is to use this energy density to develop coarse-grained expressions for the stress in the network and its elastic properties. For this purpose, and in analogy with the tensor derivatives in (1) and (3), we define the intrinsic strain derivatives

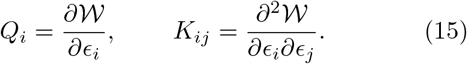

Both of these variables have the units of the energy density 𝒲, *i.e*. N/m. To physically interpret these quantities, let’s assume for the moment that the springs each respond independently and linearly to strain. Then, it is straightforward to show that 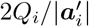 is the magnitude of the tension of spring *i*, divided by the triangle’s resting area; and 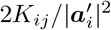are the entries of a diagonal matrix collecting the spring constants divided by the triangle’s resting area. Note that, since the triangle’s resting area is 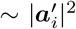, we have that *Q*_*i*_ is proportional (with a geometrical prefactor arising from the specific shape of the triangle) to the tension of the deformed spring divided by its resting length, and *K*_*ij*_ = *K*_*ji*_ represents, in general, six independent nominal rigidities. This interpretation can be used to define the conjugate quantities, *q*_*i*_, *k*_*ij*_, that apply to lengths and areas in the deformed configuration,

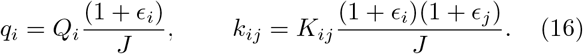

We will refer to *q*_*i*_, *k*_*ij*_, as the spatial tension densities and spatial rigidities of the unit triangle.

Using Eqs. (16), the definitions of stress tensor (1), (4), the definitions of elasticity tensors (3), (6), and the identities in (14), we obtain

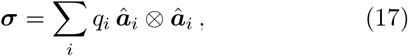

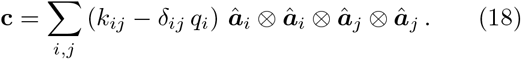

For later use, we note that the relationship (17) and the fact that each ***â***_*i*_ has unit length imply that the trace of the Cauchy stress tensor is equal to the sum of the spatial tension densities,

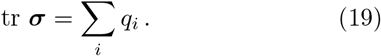

Now that we have defined continuum properties of the deformed network, we can formulate its incremental stress-strain relationship (8), and identify the effective elastic moduli of the network. In particular, with (17) and (18) we can calculate the elements of the two elasticity tensors ***m*** and ***γ*** in terms of the parameters of the unit triangle. Note that the spatial tension densities *q*_*i*_ appear in both the network’s Cauchy stress (17) and in the spatial elasticity tensor (18). This tension dependence will be explicit in ***γ***, and in a distinctive term of ***m***. Since the tensions are zero in the resting configuration, these terms collectively represent the geometric stress-stiffening of the network.

To simplify the expressions and to identify effective elastic moduli, we will utilize from hereon the Voigt notation that is commonly used in engineering, where the stress and strain elements are represented in column arrays and the elasticities as stiffness matrices. For example, in our two-dimensional system, the Cauchy stress in (17) reads

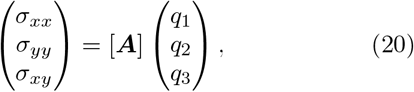

where

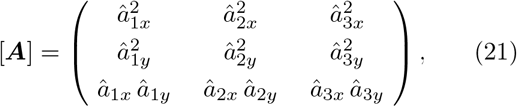

and *â*_*ix*_, *â*_*iy*_, are the two vector components of ***â***_*i*_, the deformed unit vector of the spring *i*. There are also three independent elements of *δ****σ***, three of *δ****ε***, and one of *δ****ω***, and we can write the stress increment (8) as

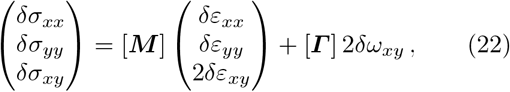

where the stiffness matrix, [***M*** ], and the rotational influence matrix, [**Γ** ], contain the elements of the tensors ***m*** and ***γ***, respectively. It is easy to show that

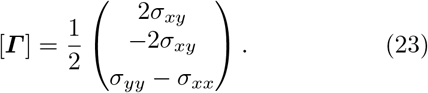

Thus, the rotation mode, *δω*_*xy*_, only contributes to equation (22) when the stress field is non-isotropic.

With some manipulation we can find

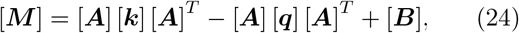

where [***k***] is a matrix with elements *k*_*ij*_, [***q***] is a diagonal matrix with elements *q*_*i*_, and [***B***] collects the Cauchy stress terms of (9),

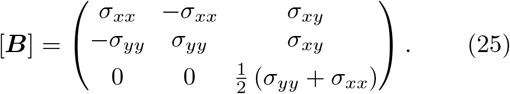

The first two terms in (24) represent the spatial elasticity tensor (18) in matrix form. We denote [***A***] [***k***] [***A***]^*T*^ ≡ [***M*** ^*k*^], the structural stiffness matrix. The remaining terms, − [***A***] [***q***] [***A***]^*T*^ +[***B***] ≡ [***M*** ^*q*^], depend on the spatial tensions *q*_*i*_ and constitute the geometric stress-stiffening part of the stiffness.

### C. Orthotropic symmetry

Large deformations strongly affect the anisotropy of materials. As an exception, for orthotropic materials that are deformed orthogonally with respect to their symmetry axis, the symmetry is preserved. This is particularly relevant for *E. coli* cell-wall mechanics (part II). In this section, we will thus specialize the results of the previous section to an orthotropic network. We will assume that the nodes of the network are torqueless hinges so that *K*_*ij*_ = *δ*_*ij*_*K*_*i*_ and the matrix [***k***] is diagonal (in part II we will justify this assumption for *E. coli* PG) with corresponding entries

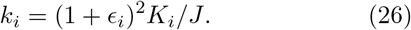

We will further assume that a prior deformation has been applied orthogonally and along the axes of symmetry. For convenience, we will set the Cartesian axes 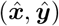 along the axes of symmetry, so that ***F*** is represented by a diagonal matrix whose entries we will denote by the stretch ratios *λ*_*x*_ and *λ*_*y*_ (and thus, *J* = *λ*_*x*_*λ*_*y*_). With these assumptions, both the relaxed and deformed configurations must be invariant to the reflections 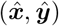 → (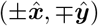), and the shear stress vanishes, *i.e. σ*_*xy*_ = 0.

Finally, we will assume, without further loss of generality, that spring *i* = 1 is parallel to the *x* axis,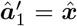, and that springs *i* = 2, 3, are mirrored segments sharing the same rigidities, *k*_3_ = *k*_2_. In this arrangement, only a single angle *ϕ* = *θ*_1_ = *θ*_3_ characterizes the shape of the triangle, or 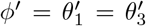 in the resting configuration. Then,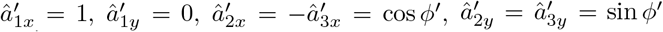 and similarly for the deformed unit vectors with angle *ϕ*. The matrix [***A***] follows from (21),

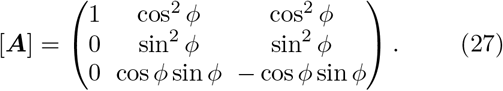

Furthermore, from (12), the two mirrored segments of the triangle must have identical strains, *ϵ*_3_ = *ϵ*_2_, and the relaxed and deformed angles are related by

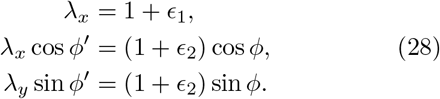

The Cauchy stress (20) reduces to

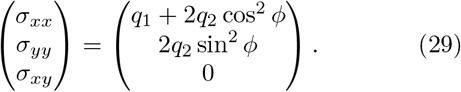

With relations (27)-(29), we can readily compute the stiffness matrix, [***M***], and the rotational influence matrix, [**Γ**]. Since *σ*_*xy*_ = 0, the latter shows, through (23), that the rotational mode *δω*_*xy*_ only affects the increment of shear stress *δσ*_*xy*_, and that only when *σ*_*yy*_ ≠ *σ*_*xx*_. Rather than writing out all the components of [***M***], we will instead interpret the stiffnesses through the standard definition of the *compliance* matrix of an orthotropic material,

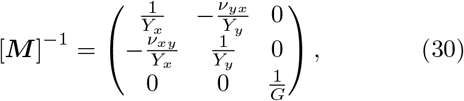

in which *Y*_*x*_ and *Y*_*y*_ are the Young’s moduli in the *x* and *y* directions, ν_*xy*_ and ν_*yx*_ are Poisson’s ratios, and *G* is the shear modulus. The increment of the shear stress then is given by the formula

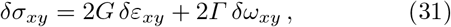

where Γ = (*σ*_*yy*_ − *σ*_*xx*_) */*2 is the rotational stiffness, the nonzero element of [**Γ**]. When the network is relaxed (so that the tension densities are zero and *λ*_*x*_ = *λ*_*y*_ = 1), *Γ* = 0 and the compliance elements take a much simpler form,

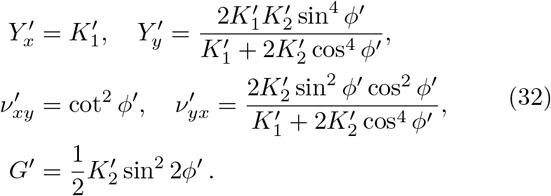

However, it is important to emphasize that a finite prestrain can substantially modify these relationships, as we will demonstrate in the next section.

## II. *E. COLI* PEPTIDOGLYCAN

We now apply the results of part I to find the minimal model for the *E. coli* PG that it is consistent with experimental data. *E. coli* is a rod-shaped bacterium. The PG layer forms an approximately spherocylindrical closed shell (sacculus), which, under normal conditions, is in a state of large deformation due to a turgor pressure of ∼ 1 atm (∼ 13% pre-strain). In this mechanical regime we assume that [***k***] is diagonal (spring network with torqueless hinges) and we will directly use the equations derived in section I C. We justify this assumption in the following.

The PG layer is a 2D-connected polymer network, made of glycan strands covalently cross-linked by peptides. Glycan strands are rigid and span distances between tens to hundreds of nanometers [29], while the peptide cross-links are flexible, with a contour length of ≈ 4 nm. Essentially, the statistical mechanics of PG is the one of a filamentous polymer network [30] where the filaments are much more rigid than the cross-links. In such a network, bending and non-affine rotations of the filaments become geometrically prohibited for large deformations [23, 30]. The deformability of the network is then largely determined by the collective response of the crosslinks [30, 31]. In other words, the elastic constants of the material are not independent, but are all coupled to the rigidity of the peptides. We can capture this situation with a triangular network with torqueless hinges, where the spring constants of the unit triangle are, in principle, different but follow the same dependence with strain. We will express this property in (34) when we introduce the response function of the springs.

We can estimate the stress in the PG network of *E. coli* and some of its mechanical properties from available experimental data. In the following, we will treat the spherical end caps of the sacculus separately from the central cylindrical region. The pre-stress of the PG layer is caused by the physiological turgor pressure, *p*, that bacteria maintain. The stress is proportional to the product of the pressure and the diameter of the sacculus, *D*, via the Laplace relationship. The cylindrical geometry of the sacculus dictates that the axial stress and the stress in the end caps equals 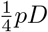, while the circumferential stress in the cylindrical region (hoop stress) equals 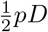, twice the axial stress, leading to a mean pre-stress equal to 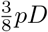. The molecular machinery elongating the cylindrical part between cell division events and the one forming the end caps during cell division are different [3], causing differences in the fine structure and mechanical properties of PG. In the cylindrical region, glycan strands have a preferred circumferential orientation [29, 32], providing this region with orthotropic symmetry. We will take *x* as the circumferential direction (hoop direction) and *y* as the axial direction. In the end caps, there is no preferential orientation of glycans, the PG is isotropic, and our choice of coordinate axes is unimportant.

Shrinking *E. coli* cells in high-osmolarity media and bringing turgor pressure to zero [13, 17, 19–22, 33] showed that the stretch ratios *λ*_*x*_, *λ*_*y*_, are similar, with values in the range 1.05–1.20. Such a uniform isotropic deformability of the sacculi under varying turgor pressures implies that the aspect ratio of the cells is preserved during osmotic challenges. This appears to be an evolved and controlled feature of the *E. coli* cell wall. The aspect ratio of *E. coli* does change during growth, when the diameter stays constant and the length increases, but at a given growth stage it is independent of the growth medium [34] although size, turgor pressure and pre-strain vary [17]. In the rest of this analysis we take the prestrain to be isotropic in all regions, so *λ*_*x*_ = *λ*_*y*_ = *λ, ϵ*_1_ = *ϵ*_2_ = *ϵ*_3_ = *ϵ* = *λ* − 1, *J* = *λ*^2^ = (1 + *ϵ*)^2^ and *q*_*i*_ = *Q*_*i*_*/*(1 + *ϵ*), *k*_*i*_ = *K*_*i*_. As a result, the angles between the springs remain equal and invariant through the deformation, *ϕ* = *ϕ*^*′*^. We will set *ϕ*^*′*^ = *π/*3, generating a network of equilateral triangles with a resting Poisson’s ratio 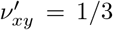, consistent with results from atomic simulations [24]. The Cauchy stress components of the deformed network are

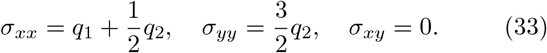

As justified earlier, the different properties of the springs are not independent but are determined by the collective response of the peptide crosslinks. Thus, the springs can exhibit different rigidities but have the same basic response to deformation. We express this generically as

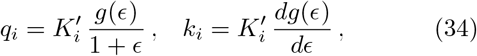

where *g*(*ϵ*) represents the shared response function. The simplest example of a response function is that of a linear spring, *g* = *ϵ* and *dg/dϵ* = 1, for which 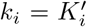 are constant at all strains. We can expect any reasonable response function to have this behavior in the limit of small strains. In section II B, we will explore the effect of intrinsic nonlinearities using the exponential strain hardening law

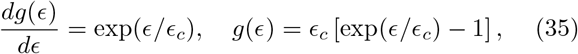

where *ϵ*_*c*_ defines the onset of the nonlinear response. This exponential model is similar to constitutive models used in other biological contexts [35]. As required, for small deformations this hardening law reduces to a linear spring model, *g* ≈ *ϵ* when *ϵ* ≪ *ϵ*_*c*_.

### A. Cylindrical region

In the cylindrical region, the orthotropic symmetry of the network demands that the rigidities of the two mirrored springs are equal, 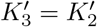 (section I C). However, 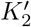 will differ from 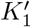to accommodate the preferred orientation of the glycan strands. The relationship between 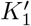 and 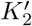 can be inferred from (33), computing the stress ratio

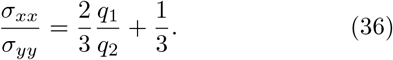

The Laplace relationship predicts that 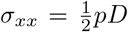 and 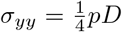, so that the stress ratio will remain equal to 2 at all levels of deformation. Thus, the ratio of forces must be *q*_1_*/q*_2_ = 5*/*2 and, due to (34), the ratio of spring rigidities must also be 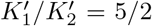. Furthermore, the expressions in (32) predict a shear modulus 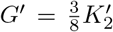 and a ratio of Young’s moduli 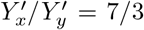, consistent with direct measurements of *E. coli* sacculi by AFM [8]. These relationships between the elastic constants reduce the description of the network to two free parameters: the response function *g*(*ϵ*) and one independent elastic constant, for example the shear modulus in the relaxed state, *G*^*′*^. It is convenient, however, to present our results in terms of the stress invariant tr ***σ*** and the following function of this invariant:

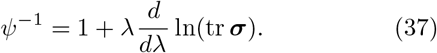

In our case, with (19) and (34), we get

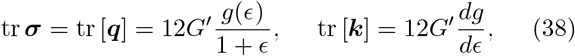

and, therefore, since *λ* = 1 + *ϵ*,

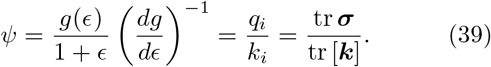

The function *ψ* represents the ratio of tension density to rigidity of the springs. It is independent of the resting rigidities and—as a consequence of the shared response function—has the same value for all springs. Therefore, *ψ* also represents the network’s overall ratio of tension to rigidity, and we will refer to it as the generalized prestrain. In particular, for small deformations the response function is linear and the generalized pre-strain is equivalent to strain, *ψ* = *ϵ/*(1 + *ϵ*) ≈ *ϵ*.

In terms of *ψ* and tr ***σ***, the stiffness matrix (24) can now be written in the form

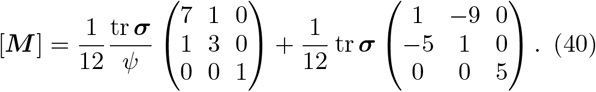

The first term represents the structural stiffness matrix of the network, [***M*** ^*k*^], and the second term is the geometric stress-stiffening contribution, [***M*** ^*q*^]. By calculating [***M***]^−1^ from (40) and comparing it with the general engineering form of the compliance matrix (30), we readily find the following relations for the elastic moduli and Poisson’s ratios:

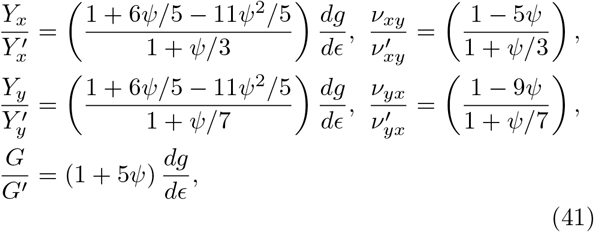

where, as in (32), the primed quantities denote the values of the resting configuration,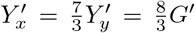 and 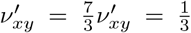. The *ψ*-dependent factors in parentheses indicate the relative effect of the geometric stress-stiffening. In the limit *ψ* → 0 the stiffness (40) is dominate by the structural stiffness matrix [***M*** ^*k*^] and, from (41), we find that the quantities 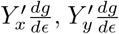 and 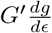 are the corresponding values of the structural moduli. Since *dg/dϵ* → 1 as *ϵ* → 0, the structural moduli reduce to the resting values in the limit of zero strain as expected.

### B. Strong strain hardening regime

The specific form of the generalized pre-strain *ψ*(*ϵ*) depends on the response function *g*(*ϵ*). For simplicity, we use the exponential strain hardening law (35), where the onset of the nonlinear response is defined by the characteristic strain *ϵ*_*c*_—but our general conclusions will not depend on the particular model chosen. For exponential strain hardening, the generalized pre-strain reads

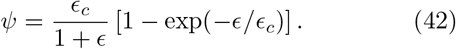

We focus our analysis on observed physiological prestrains of *E. coli, ϵ* ≈ 0.13 ± 0.08 [13, 17, 19–22, 33]. Depending on the relative value of the strain *ϵ/ϵ*_*c*_, the generalized pre-strain shows two distinct regimes, evident in Fig. 2A where *ψ*(*ϵ*) is plotted using different values of *ϵ*_*c*_. For *ϵ/ϵ*_*c*_ ≪ 1 the response of the material is linear, 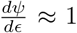, tr ***σ*** ≈ 12*G*^*′*^*ϵ* and tr ***σ****/ψ* ≈ 12*G*^*′*^. For *ϵ/ϵ*_*c*_ ≫1 the material is strongly strain hardening, leading to a plateau in *ψ*(*ϵ*). Indeed, a strong hardening regime is generally characterized by 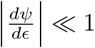, regardless of the specific model. In this regime, the definition (37) can be written as 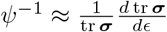 and, therefore,

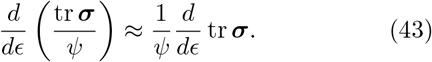

**FIG. 2:**
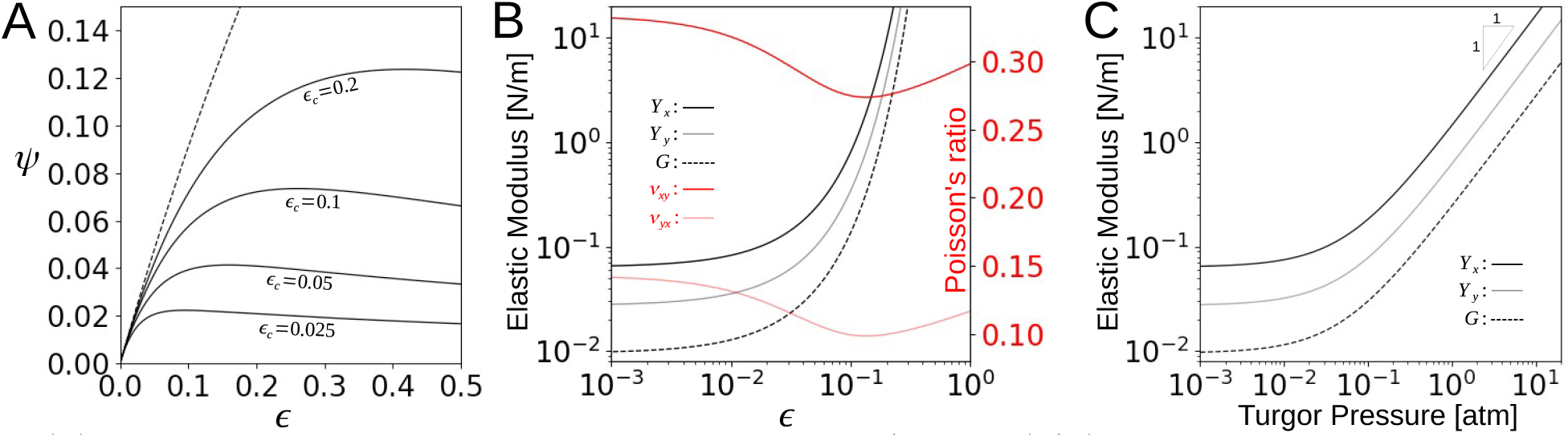
(A) *ψ* as a function of *ϵ*, as predicted by the hardening law *dg/dϵ* = exp (*ϵ/ϵ*_*c*_), for different *ϵ*_*c*_. Dashed line represents the linear spring model *ψ* = *ϵ/*(1 + *ϵ*). (B) Compliance coefficients as a function of the strain for *ϵ*_*c*_ = 0.04 (exponential hardening) and *G*^*′*^ = 10^−2^N/m. (C) Elastic moduli, same as in (B), plotted as a function of the turgor pressure of the cells.

This universal behavior implies that, to leading order, the stiffness matrix increases linearly with the stress through both terms in (40). Since the Laplace relationship states tr ***σ*** ∼ *pD*, the elements of [***M***] can be taken as linearly dependent on turgor. This is evident in Fig. 2C, where the elastic moduli are plotted against the turgor pressure.

Linear relationships between elastic constants and turgor have previously been postulated in the literature to model the stress-stiffening response of *E. coli* PG [11, 18]. The postulate in [11, 18] was based on the theoretical analysis of Boulbitch *et al*. [23], where the elastic constant *m*_*yyyy*_ of PG is shown to exhibit the asymptotic limit 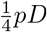. Interestingly, there appear to be two ways in which this linear relationship can be reached. In contrast to the strong strain hardening postulated in the current work, Boulbitch *et al*. [23] describe a softening regime for extreme strains. Since the trace of the stress tensor at the cylindrical region is tr 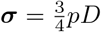, our stiffness matrix gives 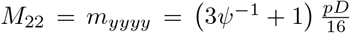, so the limit 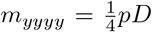 requires that *ψ* = 1. However, this value for *ψ* would imply that the spatial elasticity in (18) becomes negligible, or equivalently, [***M*** ] = [***B***] in (24). In other words, the Young’s moduli vanish, *Y*_*x*_ = *Y*_*y*_ = 0, as predicted by (41). This limit appears to be nonphysiological. In the next section, we will estimate *ψ* from experimental results. The analyzed data is consistent with the PG operating in the strong strain hardening regime.

### C. The generalized strain *ψ* of *E. coli*

An estimate of *ψ* requires measurements of cell wall deformations under different turgor pressures. There are no reported direct measurements of the relation between pre-strain and turgor. Indirect results by Cayley *et al*. [17], however, indicate a highly non-linear dependency of the cell size on turgor. The authors estimated a population average volume of *V*_0_ = 1.4 fL for unpressurized *E. coli* cells (independent of growth conditions), of *V* = 2.0 fL at a turgor pressure *p* = 1.5 ± 0.3 atm in a growth medium of low osmolality of 0.10 Osm, and of *V* = 2.2 fL at a turgor pressure *p* = 3.1 ± 0.4 atm in a growth medium of very low osmolality of 0.03 Osm. From these results we can estimate the parameters of our spring network model. The stretch ratio is *λ* = (*V/V*_0_)^1*/*3^. To obtain the pressure dependent diameter of the cells, we use the observation that the population average value of the aspect ratio of *E. coli* is approximately 4.0 and independent of the pressure [34]. Using the spherocylinder model for the shape, we find that *D* ≈ 0.70 × *V* ^1*/*3^. From the results of Cayley *et al*. [17], the Laplace relationship, tr 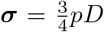, and the relationships of diameter and volume, we therefore estimate tr ***σ*** ≈ 0.10 N/m and tr ***σ*** ≈ 0.21 N/m for *λ* = 1.13 and *λ* = 1.16, respectively. On a log scale this implies an increment Δ ln(tr ***σ***) ≈ 0.74 for a stretch ratio difference Δ*λ* ≈ 0.03. Thus, from (37) we obtain a generalized pre-strain *ψ* ≈ 0.034 for *ϵ* ≈ 0.15.

The fact that *ψ* is significantly smaller than *ϵ* implies a highly non-linear response, because otherwise *ψ* would be approximately equal to *ϵ*, as discussed earlier. Interpreting the estimates for *ψ* with the exponential strain hardening model (42) shows that *ϵ*_*c*_ ≈ 0.04 and the generalized pre-strain remains relatively constant at *ψ* = 0.03 ± 0.004 for the entire physiological range of strain 0.05 *< ϵ <* 0.20. Using (38) we can also estimate the resting shear modulus *G*^*′*^ ≈ 0^−2^ N/m. We insert these values into the expressions (41) to compute the strain-dependent elastic moduli for the bacterial cell wall (Fig. 2B). The Young’s moduli increase rapidly for *ϵ >* 0.01, while the Poisson’s ratios remain approximately constant. The narrow range of variation we found for *ψ* indicates that the PG network always operates in the strong strain hardening regime for physiological conditions, such that the stiffness matrix is proportional to tr ***σ***. Consequently, all elastic moduli are proportional to stress, and scale linearly with the pressure: *Y*_*x*_ ≈ 13.8 *pD, Y*_*y*_ ≈ 5.92 *pD* and *G* ≈ 2.34 *pD* (Fig. 2C).

Because *ψ* ≪ 1, the nonlinearity associated to the collective response of the peptide crosslinks, expressed through *g*(*ϵ*) in (34), dominates over the geometric stress stiffening. Geometric stress stiffening only accounts for about 15% of the shear modulus, *G* ≈ 1.15 *G*^*′*^*dg/dϵ* (Eqs. 41), and makes an even smaller contribution to the Young’s moduli,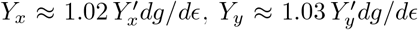. Poisson’s ratios are ν_*xy*_ ≈ 0.28 and ν_*yx*_ ≈ 0.10. The magnitude of the rotational stiffness in (31), which only arises due to geometric stress-stiffening, is only about five percent of the shear modulus,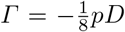. In summary, it is the intrinsic strain hardening of the peptide crosslinks, rather than geometric stress stiffening that is primarily responsible for the observed stiffening of the turgid *E. coli* cell wall.

### D. Spherical end caps

In the spherical end caps of the sacculus, no orientational order is observed in the glycans. Thus, we can directly use the isotropic condition 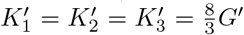 to obtain

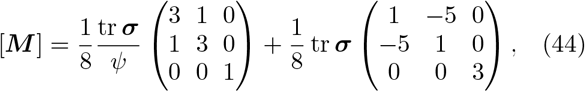

where tr ***σ****/ψ* = tr [***k***] = 8*G*^*′*^*dg/dϵ*, and *ψ*(*ϵ*) and *g*(*ϵ*) are defined as before in (37) and (34). In the end caps, the stress invariant, tr 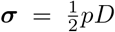, is two-thirds of the stress invariant in the cylindrical region,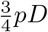. This implies a structural shear modulus 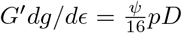. Interestingly, this relationship also applies to the cylindrical region, so the value of the resting shear modulus we estimated previously, *G*^*′*^ ≈ 10^−2^N/m, is valid in the whole sacculus. However, the stiffness matrix (44) describes different strain-dependent elastic moduli and Poisson’s ratios from those in the cylindrical region,

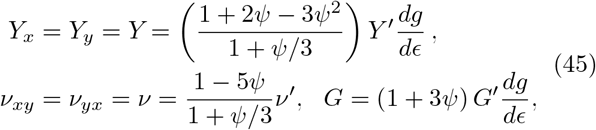

where 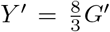. We can again use *ψ* = 0.03 ± 0.004, and estimate the turgor-pressure dependence of the elastic moduli in the end caps: *Y* ≈ 1.05 *Y* ^*′*^*dg/dϵ* ≈ 5.69 *pD*, ν ≈ 0.28 and *G* ≈ 1.09 *G*^*′*^*dg/dϵ* ≈ 2.21 *pD*. Because of the isotropic stress, equation (23) shows that the rotational stiffness, which is purely geometrical, vanishes at the spherical regions.

### E. Discussion

We have here developed and analyzed a mechanical model of the *E. coli* PG layer, which dominates the elastic response of the bacterial cell wall, using a 2D regular triangular network. In our analytical approach, we have built a bridge to an elastic-continuum description and present a self-contained method to study stress-strain relationships with an emphasis on large deformations and arbitrary anisotropies. We inserted parameters into our model based on AFM imaging of peptidoglycan layers [29, 32] and measurements of the passive response of *E. coli* to osmotic challenges [17, 19–21]. An important assumption of the model reflects the observation that the aspect ratio of *E. coli* is not affected by the magnitude of the turgor pressure [19–21, 34]. This has allowed us to characterize the PG network with an isotropic prestrain, which greatly simplified the analysis. A crucial result of our analysis is that the *E. coli* PG is in a state of strong strain hardening under physiological turgor pressures, leading to a stiffness matrix that is linearly proportional to the turgor pressure.

Although these results hold without a specific identification of the springs of our model with the actual molecular components of the PG layer, this study opens up a path toward a molecular-scale interpretation. The diagonal form we have used for [***k***] to compute the stiffness matrix (24) is based on the idea that the observed deformations of the PG layer (and, therefore, its elastic energy) mainly reflect the stretching of the flexible peptide crosslinks. While rigid glycan filaments store relatively small amounts of elastic energy they do impose, because of their length, crucial geometric constraints. The mechanical properties of the springs of our model include this effect indirectly, as explained in the following.

Individual peptide cross-links can be modeled as non-linear springs that behave as linear entropic springs, with spring constant *k*_*sp*_ ≈ 2.3 × 10^−2^ N/m [6], up to strain values of about 1. The onset of strain hardening of individual peptides is thus well above the largest overall pre-strains observed in the *E. coli* cell wall, of *ϵ* ≈ 0.20. Why, then, does the PG layer strain harden under physiological conditions? The answer lies in the constraints imposed by the glycan strands. If we were to ignore these constraints, *k*_*sp*_ would define the magnitude of the shear modulus of the whole network. Using realistic numbers, the structural shear modulus of a hexagonal network made up of only peptide crosslinkers, without glycans, would be 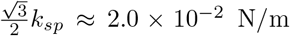 This value is indeed close to the resting shear modulus *G*^*′*^ = 10^−2^ N/m, which we estimate for *E. coli* sacculi, but is two orders of magnitude smaller than the structural shear modulus of the pressurized shells *G*^*′*^ exp (*ϵ/ϵ*_*c*_) ≈ 1.5 N/m, for *ϵ* = 0.20 and for *ϵ*_*c*_ = 0.04. The long glycan strands, to which the peptide bridges are linked, enforce a broad distribution of peptide strains with a heavy tail extending to strain values much larger than the macroscopic average strain of the cell. This is the mechanism behind the dramatic strain hardening effect observed (Fig. 2). Some aspects of this general phenomenon have been explored with an *affine* elastic theory of rods embedded on a continuum space of flexible crosslinkers [30, 31].

In our regular-network approximation, we capture the experimentally determined response of the real PG on a mesoscopic scale by defining two types of springs to allow for anisotropy, and by shifting the onset of nonlinearity of the springs to a lower strain value than the one expected for the peptide crosslinks. This approach takes the tail of the strain distribution in the real PG into account. The challenge for future work is to explain the overall mechanical behavior of the bacterial cell wall on the basis of the structural statistics of the real PG. Our multi-scale modeling approach will enable more detailed investigations of other extremal biological materials in general. In bacteria, in particular, it will make it possible to quantitatively understand the response of the cell wall to antibiotics that interfere with cell wall synthesis.

## F. Acknowledgments

We dedicate this article to Professor Alex Levine, who recently passed away. Intensive discussions with Alex started our work on this topic. This material is based upon work supported by the National Science Foundation under Grant Nos. 2221771 and 2221772.

## Notes

### Competing Interest Statement

The authors have declared no competing interest.

